# Andes virus genome mutations that are likely associated with animal-model attenuation and human person-to-person transmission

**DOI:** 10.1101/2023.01.18.524667

**Authors:** Carla M. Bellomo, Daniel O. Alonso, Unai Pérez-Sautu, Karla Prieto, Sebastian Kehl, Rocio M. Coelho, Natalia Periolo, Nicholas Di Paola, Natalia Ferressini-Gerpe, Jens H. Kuhn, Mariano Sanchez-Lockhart, Gustavo Palacios, Valeria P. Martínez

**Author notes:** These authors contributed equally to the study (Carla M. Bellomo, Daniel O. Alonso, Gustavo Palacios, Valeria P. Martínez). Address correspondence to Gustavo Palacios, or Valeria P. Martínez,.

## Abstract

**Abstract:** We performed whole-genome sequencing with bait-enrichment techniques to analyze Andes virus (ANDV), a cause of human hantavirus pulmonary syndrome. We used cryopreserved lung tissues from a naturally infected long-tailed colilargo; early, intermediate, and late cell-culture passages of an ANDV isolate from that animal; and lung tissues from golden hamsters experimentally exposed to that ANDV isolate. The resulting complete genome sequences were subjected to detailed comparative genomic analysis against American orthohantaviruses. We identified four amino-acid substitutions related to cell-culture adaptation that resulted in attenuation of ANDV in the typically lethal golden hamster animal model of hantavirus pulmonary syndrome. Mutations in the ANDV nucleocapsid protein, glycoprotein, and small nonstructural protein open reading frames correlated with mutations typical for ANDV strains associated with increased pathogenesis in the small animal model. Finally, we identified three amino-acid substitutions, two in the small nonstructural protein and one in the glycoprotein, that were only present in the clade of viruses associated with person-to-person efficient transmission. Our results indicate that there are virulence-associated and transmission-associated single-nucleotide polymorphisms that could be used to predict strain-specific ANDV virulence and/or transmissibility.

**Importance:** Several orthohantaviruses cause the zoonotic disease hantavirus pulmonary syndrome (HPS) in the Americas. Among them, HPS caused by Andes virus (ANDV) is of great public-health concern because it is associated with the highest case-fatality rate (up to 50%). ANDV is also the only orthohantavirus associated with relatively robust evidence of person-to-person transmission. This work reveals nucleotide changes in the ANDV genome that are associated with virulence attenuation in an animal model and increased transmissibility in humans. These findings may pave the way to early severity predictions in future ANDV-caused HPS outbreaks.

## INTRODUCTION

Approximately 25 rodent-borne orthohantaviruses (*Bunyavirales: Hantaviridae: Orthohantavirus*) have been identified as etiologic agents of human hantavirus pulmonary syndrome (HPS) in the Americas (1). In Argentina, most HPS cases are caused by Andes virus (ANDV) and somewhat uncharacterized ANDV-like viruses (e.g., Buenos Aires virus [BASV], Lechiguanas virus [LECV], Orán virus [ORNV]). HPS has a case-fatality range of 21–50%, with ANDV typically causing the highest lethality (2–5). American orthohantaviruses are pathogenic for humans and subclinically infect cricetid rodents in nature; ANDV is primarily maintained by long-tailed colilargos (*Oligoryzomys longicaudatus* (Bennett, 1832)) (6).

The route of orthohantavirus transmission to humans is typically zoonotic, i.e., *ex vivo* without intermediate vectors (7). However, in 1996, an HPS outbreak caused by ANDV strain Epilink/96 that began in El Bolsón, Río Negro Province, Argentina, was attributed for the first time to person-to-person transmission (4, 8, 9). Sporadic HPS outbreaks with very limited person-to-person ANDV transmission have occurred over the last 25 years (2, 3, 10). Recently, state-of-the-art molecular epidemiology applied to a 2018–2019 HPS outbreak in Epuyén, Chubut Province, Argentina, confirmed the unique capacity of some strains of ANDV (in this instance, ANDV/Epuyén/18-19) to sustain forward orthohantavirus transmission in humans (11).

ANDV and Maporal virus (MAPV) are the only orthohantaviruses that have been documented to reproduce key features of HPS and cause lethal disease in a rodent model, i.e., golden hamsters (*Mesocricetus auratus* (Waterhouse, 1839)) (12–14). Immunocompetent golden hamsters provide uniformly lethal results when exposed to the Chilean strain ANDV/CHI-9717869 (isolated from a long-tailed colilargo collected from Lago Atravesado, Coyhaique, Aysen Region, Chile, in 1997) (12, 15) or the Argentinean strain ANDV/ARG (isolated from a long-tailed colilargo collected in the vicinity of the primordial site of discovery of ANDV [El Bolsón] in 2000) (14). However, the golden hamster model did not produce lethal results when exposed to a closely related strain, ANDV/CHI-7913 (isolated from clinical samples from a fatal case that was a family contact of the index case of an outbreak near Santiago, Chile, in 1999) (16). These findings indicated that subtle strain-specific genomic differences may have dramatic phenotypic consequences (17).

Cell-culture passaging has been associated with viral virulence attenuation for multiple orthohantaviruses in animal models (18, 19). We therefore hypothesized that serial cell-culture passaging of an ANDV known to be uniformly lethal in golden hamsters would result in attenuation, that attenuation would be traceable to specific mutations in the ANDV genome, and that these mutations may be catalysts for ANDV adaptation and therefore possible predictive markers for virulence and/or transmissibility.

## RESULTS

### Cell-culture passaging of Andes virus strain ARG results in virulence attenuation *in vivo*

Andes virus strain ARG (ANDV/ARG) is one of a select few available strains isolated directly from the rodent reservoir, long-tailed colilargos. To our knowledge, it is also the only ANDV strain directly sequenced from rodent material (passage 0 [p0]). We hypothesized that cell-culture passaging attenuates ANDV/ARG. To test this hypothesis, we passaged ANDV/ARG p9, described previously as causing 100% lethality in golden hamsters at 10 d after exposure (14), an additional 10 times in grivet Vero E6 cells (to p19). In a side-by-side comparison, all golden hamsters exposed via intramuscular injection of ANDV/ARG p9 uniformly reached euthanasia criteria, as expected, whereas 33.3% of those exposed to ANDV/ARG p19 recovered, and mock-exposed control animals uniformly survived (**Fig. 1**). Kaplan–Meier comparison of survival curves and log-rank tests (Mantel–Cox [*p*-value <0.0001, Chi-square 18.47], trend variation with the number of passages [*p*-value 0.0101, Chi-square 6.610], and Gehan–Breslow–Wilcoxon [*p*-value 0.0005, Chi-square 15.13]) demonstrated that these survival differences are statistically significant. ANDV/ARG RNA was consistently detected in golden hamster lung samples (3.8×10^6^ to 1.7×10^10^ RNA copies per 100 mg of perfused tissue) in the ANDV/ARG p9 and p19 cohorts but not in the mock-exposed control cohort.

**Fig 1.**
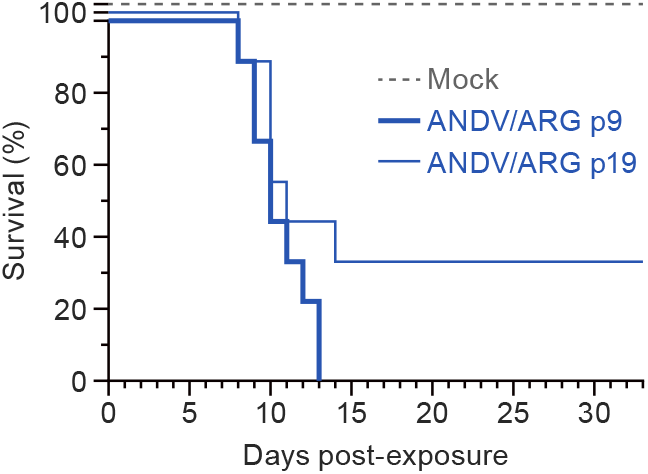
Cell-culture passaging of Andes virus results in virulence attenuation *in vivo*. Shown are Kaplan–Meier survival curves of golden hamsters inoculated intramuscularly with three different preparations until the study endpoint. ANDV/ARG, Andes virus strain ARG; p, passage.

### Phylogenetic analysis informs the evolutionary history of Buenos Aires virus and Andes virus strain ARG

We performed phylogenetic analysis of ANDV/ARG and Buenos Aires virus (BASV) small (S) and medium (M) genomic segments as well as the ANDV/ARG large (L) segment. Complete coding nucleic-acid sequences determined in this study were assessed together with previously determined sequences of ANDV, ANDV-like viruses Lechiguanas virus (LECV) and Orán virus (ORNV), BASV/BA02-C1S, and several New World orthohantaviruses (Laguna Negra virus [LANV], Sin Nombre virus [SNV], Maporal virus [MAPV], Rio Mamoré virus [RIOMV], and Choclo virus [CHOV]). Four distinct ANDV clades are apparent in the most divergent S segment tree (**Fig. 2C; details about the strains are listed in Table S1**):

1. ANDV/CHI-7913 (Chile; long-tailed colilargo) and ANDV/NRC-4/18 (Argentina; human)
2. ANDV/Epilink/96, ANDV/Epuyén/18-19, ANDV/NRC-2/97, and ANDV/NRC-6/18 (Argentina; human; all associated with person-to-person transmission);
3. ANDV/ARG (Argentina; long-tailed colilargo); and
4. ANDV/CHI-9717869 (Chile; long-tailed colilargo).

**Fig 2.**
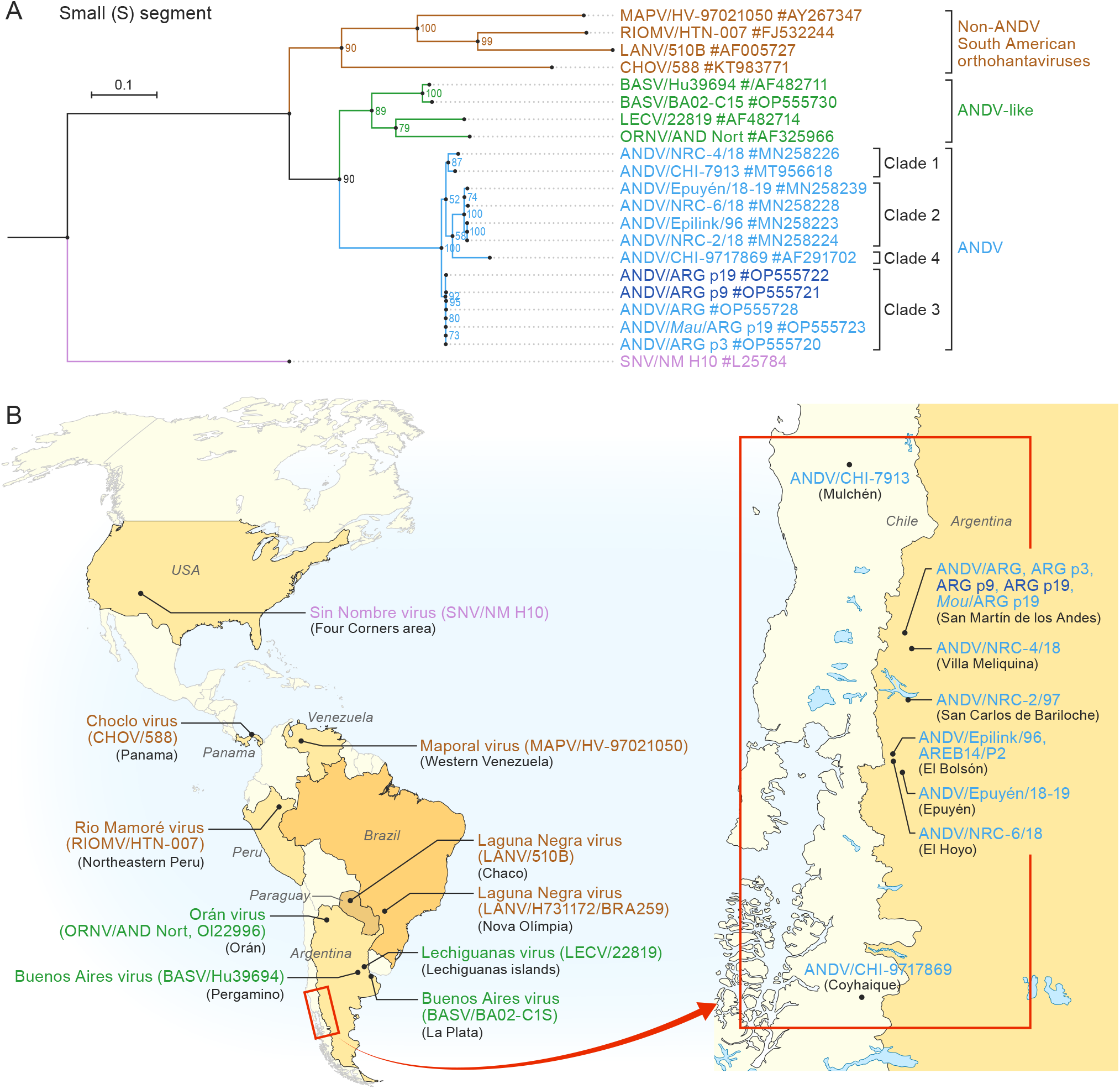
Phylogenetic analysis informs the evolutionary history of Buenos Aires virus (BASV) and Andes virus strain ARG (ANDV/ARG). (A) Small (S) segment analysis; large (L) segment and medium (M) segment analysis are included in **Figure S1**. All variants are listed with the strain name, region of origin, year of the isolation and accession number. Different colors are used for identification (brown for non-ANDV South American orthohantaviruses; green for ANDV-like viruses; light blue for ANDV strains in clades 1, 2, 4, and some in 3; and dark blue for passaged strains in clade 3). Detailed information of epidemiological history of the strains is listed in **Table S1**. (B) Geographic distribution of American orthohantaviruses strains analyzed in A. Mulchén and Coyhaique are in Chile, the other locations are in Argentina. The inset shows the endemic area of ANDV in Argentina and Chile.

ANDV/ARG is therefore not directly related to the other ANDV strains associated with person-to-person transmission. Interestingly, BASV clusters separately from ANDV *sensu stricto* together with LECV and ORNV; and ANDV/CHI-9717869 appears to be the ancestral to the ANDV species. Further, the analysis also shows that ANDV/ARG genetic distances to other strains reflect their geographic distribution (**Fig. 2D**).

### Sequencing of passaged variants of Andes virus strain ARG reveals sites of adaptation associated with attenuation in the golden hamster model

To identify genotypic differences associated with golden hamster model outcome phenotype, we sequenced the S, M, and L genomic segments of ANDV/ARG p0 (sampled from the long-tailed colilargo). The resulting isolate was seeded in Vero E6 cells for analysis of p3, p9 (14), and p19, as well as lung-tissue homogenate from golden hamsters exposed to ANDV/ARG p9. We also included a human blood sample of Buenos Aires virus (BASV/BA02-C1S) from a hantavirus pulmonary syndrome (HPS) case in La Plata, Provincia de Buenos Aires, in 2002. We obtained complete genomic sequences for all segments (>98.3% coverage) for all ANDV strains, except for the L segment from the p0 strain (46.1% coverage). Sequences are available in GenBank, under accession numbers OP555720–OPG555735.

The comparative analysis revealed only a few nucleotide changes over passages (**Table 1**), and p0 and p3 sequences were identical. By p9, three single-nucleotide polymorphisms (SNPs) were observed: two in the *S* segment (S46N in the nucleocapsid [*N*] open reading frame [ORF] and V20I in the small nonstructural protein [*NSs*] ORF) and one in the L segment (I1295M in the large protein [*L*] ORF). By p19, three additional SNPs were observed: one in the S segment (A21T), one in the M segment (S97P), and one in the L segment (P1675S); also, one reversion was observed in the S segment (affecting S46 in the *N* ORF and V20 in the *NSs* ORF). As expected, the p19 sequence had the highest number of non-synonymous substitutions. The changes were predominantly transitions (87.5%). After correction by segment length, it is evident that most nucleotide substitutions accumulated in the S segment. Surprisingly, very few SNPs were observed in the M segment. Interestingly, no reversions were detected in the genomic sequences of ANDV/ARG p9 in the lungs of golden hamsters exposed to ANDV/ARG p9. (*Note:* No data was collected from the lungs of golden hamsters exposed to ANDV/ARG p19.)

**Table 1.**
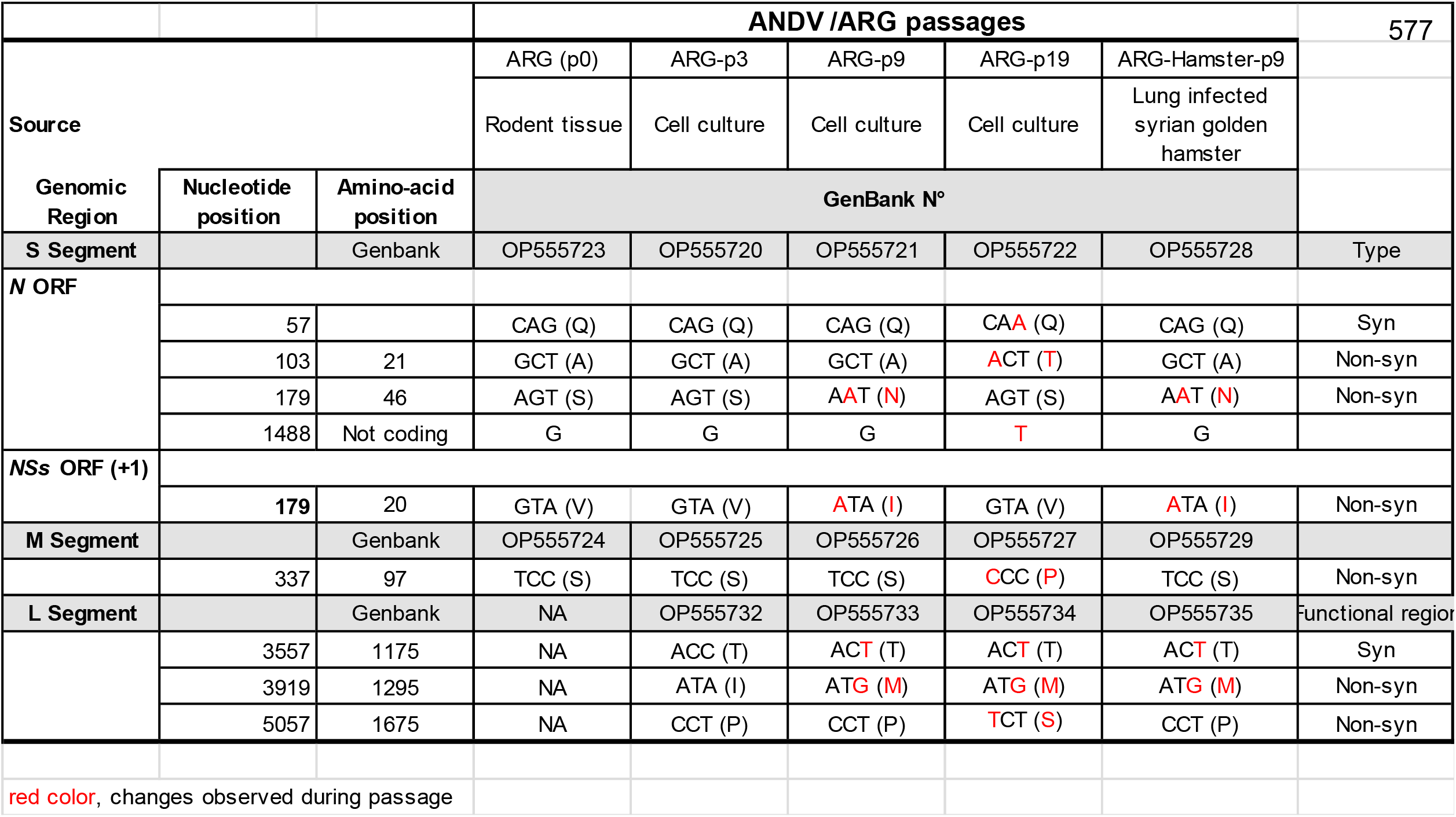
Sequencing of passaged variants of ANDV/ARG reveals sites of adaptation associated with attenuation in the golden hamster model.

|  |  |  | ANDV /ARG passages |  |  |  |  | 577 |
| --- | --- | --- | --- | --- | --- | --- | --- | --- |
| Source |  |  | ARG (p0) | ARG-p3 | ARG-p9 | ARG-p19 | ARG-Hamster-p9 |  |
|  |  |  | Rodent tissue | Cell culture | Cell culture | Cell culture | Lung infected syrian golden hamster |  |
| Genomic Region | Nucleotide position | Amino-acid position | GenBank N° |  |  |  |  |  |
| S Segment |  | Genbank | OP555723 | OP555720 | OP555721 | OP555722 | OP555728 | Type |
| N ORF |  |  |  |  |  |  |  |  |
|  | 57 |  | CAG (Q) | CAG (Q) | CAG (Q) | CAA (Q) | CAG (Q) | Syn |
|  | 103 | 21 | GCT (A) | GCT (A) | GCT (A) | ACT (T) | GCT (A) | Non-syn |
|  | 179 | 46 | AGT (S) | AGT (S) | AAT (N) | AGT (S) | AAT (N) | Non-syn |
|  | 1488 | Not coding | G | G | G | T | G |  |
| NSs ORF (+1) |  |  |  |  |  |  |  |  |
|  | 179 | 20 | GTA (V) | GTA (V) | ATA (I) | GTA (V) | ATA (I) | Non-syn |
| M Segment |  | Genbank | OP555724 | OP555725 | OP555726 | OP555727 | OP555729 |  |
|  | 337 | 97 | TCC (S) | TCC (S) | TCC (S) | CCC (P) | TCC (S) | Non-syn |
| L Segment |  | Genbank | NA | OP555732 | OP555733 | OP555734 | OP555735 | Functional region |
|  | 3557 | 1175 | NA | ACC (T) | ACT (T) | ACT (T) | ACT (T) | Syn |
|  | 3919 | 1295 | NA | ATA (I) | ATG (M) | ATG (M) | ATG (M) | Non-syn |
|  | 5057 | 1675 | NA | CCT (P) | CCT (P) | TCT (S) | CCT (P) | Non-syn |
| red color, changes observed during passage |  |  |  |  |  |  |  |  |

### Sequencing of Andes virus strain ARG reveals virulence markers when compared with pathogenic and non-pathogenic strains of Andes virus utilized in the golden hamster model

To identify potential genotypic virulence markers in the ANDV/ARG genome, we initially focused on 23 specific SNPs that had been described between the golden hamster attenuated ANDV/CHI-7913 compared to golden hamster lethal ANDV/CHI-9717869 (17). We also mapped 5 additional SNPs between those genomes, as the *NSs* ORF was not included in the original comparison (17). In 23 of those 28 positions, ANDV/ARG p0 shared nucleotide bases with attenuated ANDV/CHI-7913. ANDV/CHI-97177869 and ANDV/ARG only shared position 11 of the *Gn* glycoprotein, position 938 of the *Gc* glycoprotein, and positions 20 and 37 of the *NSs* ORFs (**Table 2**). ANDV/ARG differ from both ANDV/CHI-7913 and ANDV/CHI-9717869 in genome position 46 of the *N* ORF.

**Table 2.**
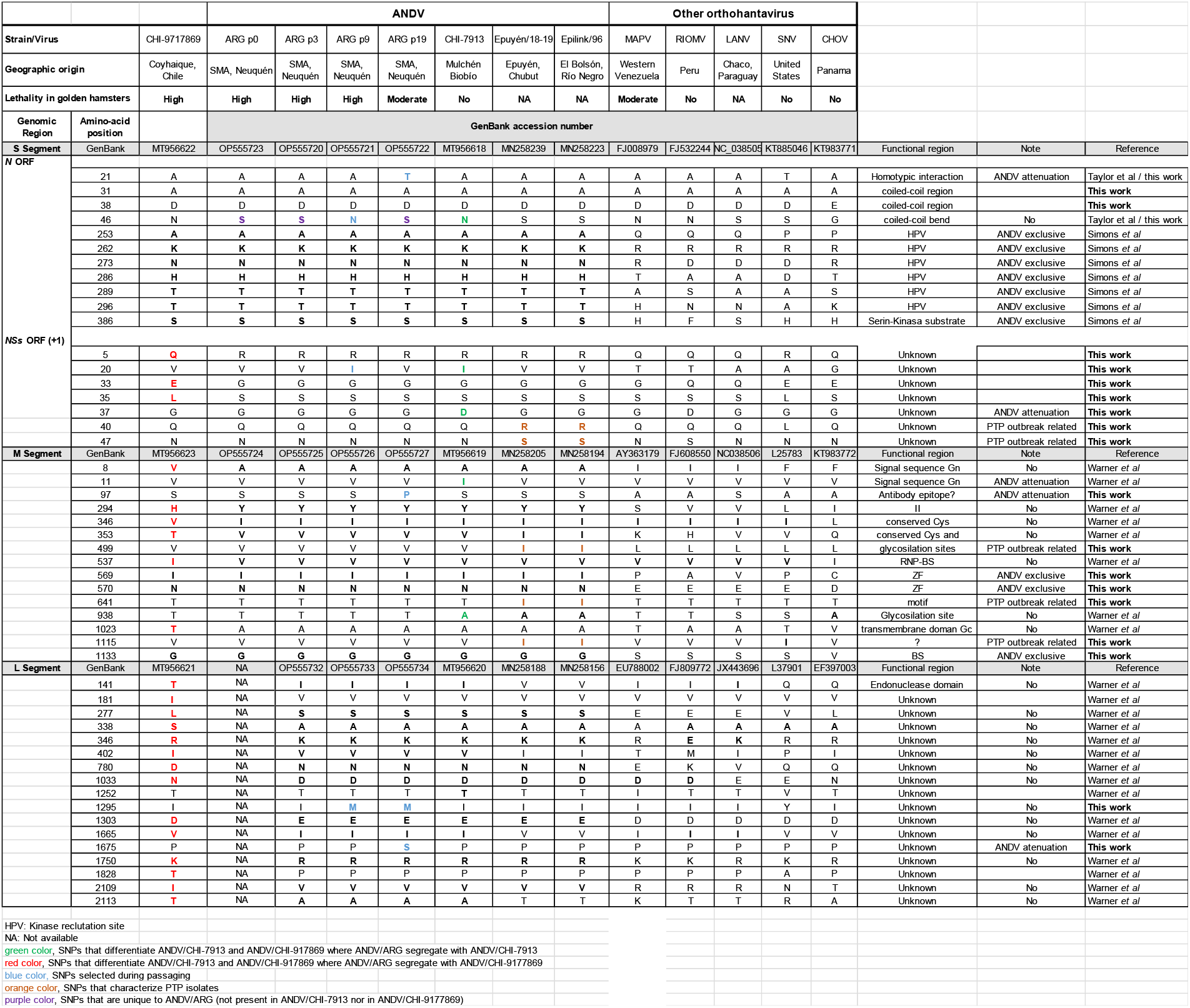
Sequencing of ANDV/ARG reveals virulence markers when compared with pathogenic and non-pathogenic strains of ANDV utilized in the golden hamster model.

Next, we focused on comparing the amino-acid changes that arose during ANDV/ARG passaging with the differences in virulence observed in the golden hamster model. We identified five: A21T in the *N* ORF, V20I in the *NSs* ORF, S97P in the *Gn* glycoprotein, and I1295M and P1675S in the *L* ORF.

T21, which only appeared in ANDV/ARG p19, occurs in a region known to participate in orthohantavirus NSs homotypic interactions (20). Additionally, we identified a second amino-acid change in the *N* ORF (S46N), which was only encoded by ANDV/ARG p9 (**Table 1 and Table 2**). Interestingly, in the same *N* ORF, Simons et al. reported an ANDV-specific kinase-recruitable hypervariable domain (HVD) in the *N* ORF by comparison of ANDV/CHI-7913 with other American orthohantaviruses and demonstrated its importance in regulating interferon (IFN) signaling (21). The HVD, which consists of 44 residues (nucleotides 252 to 296), encodes 6 characteristic amino acids (at positions A253, K262, N273, H286, T289, and T296) that are determinants of the phosphorylation of S386 in the *N* ORF, which is posited as a virulence determinant (21). Although we confirmed that S386 and five of the six residues are conserved among all ANDV and ANDV-like viruses (**Table 2 and 3**) A253 is exclusive for ANDV and has been mutated to P (BASV and LECV) or L (ORNV) in ANDV-like viruses.

**Table 3.**
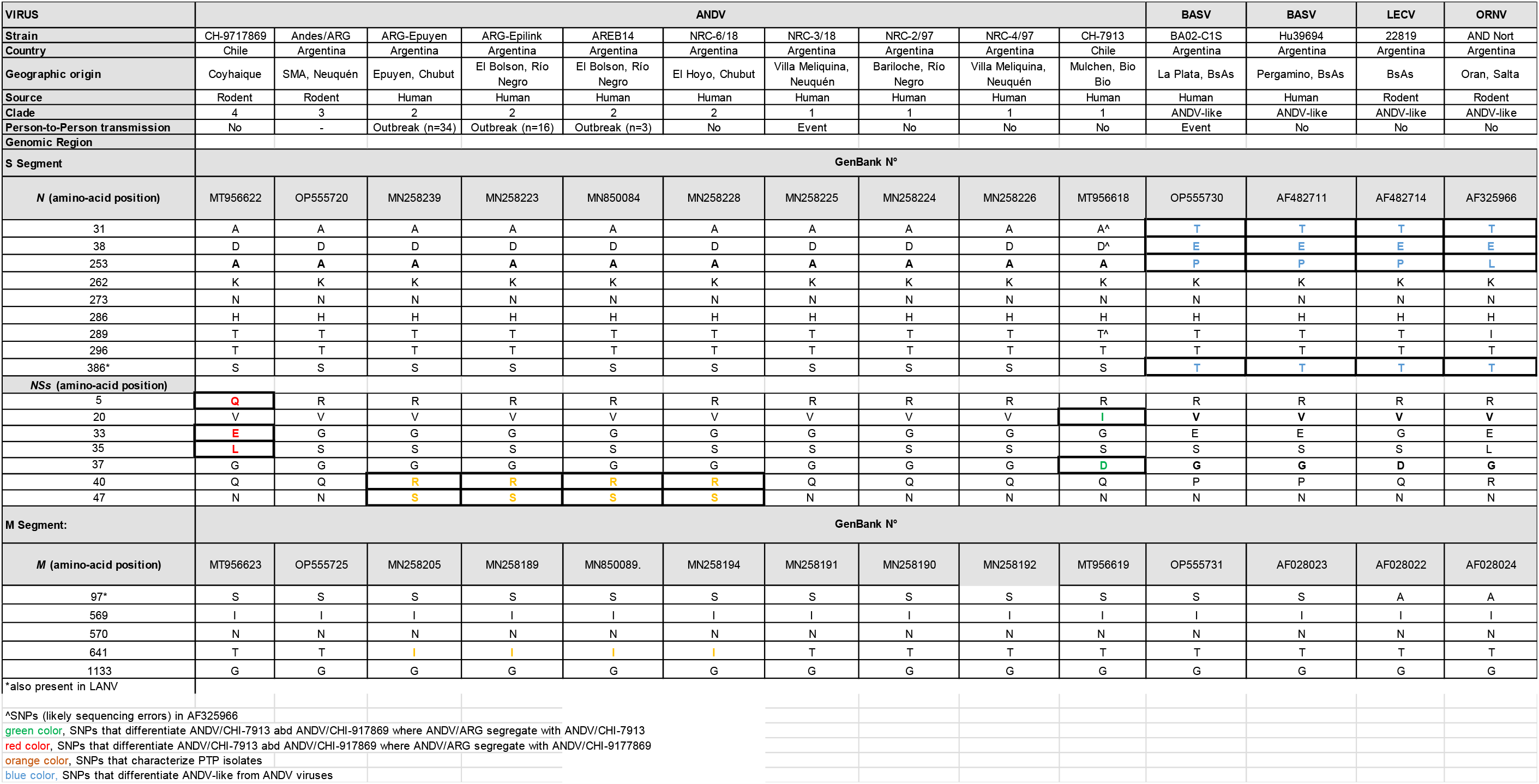
Sequencing of ANDV/ARG reveals potential transmissibility markers when comparing ANDV strains with differences of efficiency in person-to-person potential.

The recently discovered ANDV NSs antagonizes the type I IFN response by inhibiting mitochondrial antiviral-signaling protein (MAVS) signaling by binding MAV without disrupting MAVS-TBK-1 (20). In the presence of ANDV NSs, the ubiquitinylation of MAVS is reduced. The V20I SNPs in the *NSs* ORF was observed arising in the ANDV/ARG p9 strain by the same mutation in nucleotide position 179 in the S segment. (*Note:* The *NSs* ORF is in position +1 compared with the *N* ORF.) ANDV/CHI-9717869 and ANDV/CHI-7913 differ in the *NSs* ORF in five amino-acid positions (5, 20, 33, 35, and 37; **Table 2 and 3**).

The S97P Gn mutation, found only in ANDV/ARG p19, could not be associated with any functional change. The site has only been reported as part of an antibody epitope (22). The S residue is conserved in BASV and LANV Gns, whereas other orthohantavirus Gns (LECV, ORNV, MAPV, RIOMV, SNV, and CHOV) have an A in that position (**Table 2 and Table 3**).

M1295 and S1675 in the *L* ORF were encoded by ANDV/ARG p9 and p19 strains, respectively, but those genomic regions could not be associated with any functionality. M1295 has not been observed previously in nature; other orthohantavirus have I1295 (MAPV, RIOMV, LANV, and CHOV) or Y1295 (SNV) (**Table 2**). S1675 has not been observed in *L* ORFs of other orthohantaviruses, and P1675 position appears entirely conserved (**Table 2**).

### Sequencing of Andes virus strain ARG reveals potential transmissibility markers when comparing Andes virus strains with differences of efficiency in person-to-person potential

The presence of outbreak-related determinants associated with person-to-person transmission was assessed by comparing genomic sequences of ANDV strains from clades clearly associated with person-to-person transmission and ANDV strains and ANDV-like viruses (BASV, LECV, and ORNV) that had not (**Table 3**). Interestingly, BASV, the most closely related ANDV-like virus (**Fig. 2A-C**), has also been implicated in secondary transmissions but with limited efficiency (23, 24).

Only one mutation in the M segment (T641I) was unique to person-to-person-associated clade 2 strains. Only one mutation in the *S* segment (A253N) was exclusively present in ANDV, whereas S386N is conserved among ANDV strains and ANDV-like viruses. Our analysis did not include the *L* segment of ANDV-like viruses because those sequences remain unavailable.

Five *M* ORF positions were unique to ANDV genomes compared with genomes of ANDV-like viruses (amino-acid positions 97, 569, 570, 641, and 1133; **Table 3**). Three (569, 570, and 1133) are shared by all ANDV strains. S97 is encoded by all ANDV strains and BASV, whereas the T641I change was only encoded by ANDV strains from the clade associated with person-to-person transmission. However, only the latter (T641I) had been mapped in the vicinity of the absolutely conserved pentapeptide WAASA cleavage site, where signal peptidases cleave Gn and Gc (25). Since position 641 maps to a region that provides a signal to cellular peptidases, this mutation might affect the cleavage’s efficiency. Signal peptides share several characteristic features determined by their amino-acid composition (26), including a tripartite architecture with a positively charged N-terminus and a hydrophobic segment that determines the strength of the signal. T641 changes from a polar non-charged amino acid (T) to a non-polar (I) amino acid.

In comparison with the bulk of described ANDV isolates, the recently discovered ANDV *NSs* ORF presents seven sites of variation. Three are unique to ANDV/CHI-9717869 (Q5, E33, and L35) and two are unique to ANDV/CHI-7913 (I20 and D37). Intriguingly, we identified two SNPs in the *NSs* ORF at positions 40 (Q40R) and 47 (N47S) that were present only in the clade 2 strains (e.g., ANDV/Epuyén/18-19 and ANDV/Epilink/96) associated with person-to-person transmission. Both *NSs* ORF changes need to be functionally evaluated for their effect on MAVS signaling.

## DISCUSSION

Passaging in cell culture, especially when involving different hosts, usually results in virus adaptation, often affecting their virulence (19, 27, 28). However, ANDV/ARG p0 and p3 genome sequences were identical, and very few mutations were accumulated in the p9, p19, and hamster strains. The two amino-acid substitutions (A21T and S46N) in the *N* ORF mapped to the intramolecular coiled coil structure in the N-terminal region (α1 and α2), an exceptionally well-conserved region implicated in antibody recognition, formation of the ribonucleoprotein complex, and genome encapsidation (29–32). Interestingly, one adaptation appears to involve a change in the novel *NSs* ORF, which has been recently related with IFN regulation. Only a single nucleotide change (T337C) was found in the *M* segment during late passaging (p19). This is unexpected since the M segment encodes for *Gn* and *Gc*, two of the most variable regions of the genome in evolutionary terms.

Interestingly, we could also correlate some of the changes with differences in pathogenicity in a small animal model. ANDV/ARG p9 is uniformly lethal in hamsters (14). However, ANDV/ARG p19 was significantly less lethal (66.4%). Compared side-by-side, the ANDV/ARG p9 and p19 only diverged in three encoded residues (A21T in N, S97P in Gn, and P1675S in RNA-dependent RNA polymerase (RdRp) codified in the S, M, and L segments, respectively). Nevertheless, based on previous knowledge of functional domains, only the change in the N had been associated with viral replication. Structural studies of the N-terminal region of SNV and ANDV demonstrated that basic residues interact with the N core to stabilize interprotomer N association and formation of ribonucleoprotein (RNP) complexes (31). The A21T change likely affects that region, which is exceptionally well-conserved among orthohantaviruses. The region is a target of the most cross-reactive antibodies against orthohantavirus, immunodominant, and proposed to have important effects regarding N polymerization, RNP complex formation, and subcellular localization of the assembly sites (29–31). We hypothesize that A21T and other changes in N (**Table 2 and Table 3**) may affect N oligomerization dynamics. The importance of this area as a potential determinant of pathogenesis might be underscored by the observed differences in the region at positions 31 (A31T) and 38 (D38E) (**Table 3**) that define ANDV-like viruses (i.e., BASV, LECV, and ORNV). The changes, all located at the bend between the two parallel coiled regions, could potentially affect the structure of the region. On the other hand, these two changes are only encoded by ANDV-like viruses, but not by LANV, MAPV, RIOMV, or SNV (**Table 2**). Thus, if these markers are associated with pathogenesis, they would act via changes in the structure and not necessarily by SNP differences. Although the T21A mutation observed in late passages of ANDV/ARG is intriguing, A21 is conserved in BASV, LECV, and ORNV, but not SNV (**Table 2**). Collectively, this could indicate that structural changes in this area could be driving virulence differences, instead of SNPs. Moreover, our analysis confirmed that the amino-acid position S386, previously posited by Simons as a determinant of virulence (21), is conserved by ANDV, ANDV-like viruses (BASV, LECV, and ORNV), and LANV. In the N HVD, all ANDV strains share the described signature six residues, which are not found in any other orthohantavirus N HVD (**Table 2**). However, only five residues are shared with the three ANDV-like viruses, while A253 seems to be an exclusive ANDV marker (**Table 3**). We therefore suggest that A253 is an ANDV-exclusive virulence determinant and that the S386 modification and the five remaining HDV residues are pathogenic determinants for all viruses currently classified in species *Andes orthohantavirus* (i.e., ANDV and ANDV-like viruses). The *N* ORF has been associated with multiple functions associated with pathogenesis and virulence. The efficiency of orthohantavirus replication is inversely proportional to the ability of infected cells to activate MxA expression (33). The MxA protein is a critical component of the antiviral state induced by type I IFN (34). In turn, MxA protein binds to N, forming an MxA–N protein complex in a yet-to-be-defined manner (35). Moreover, the N protein also has a role in regulating the antiviral state. For instance, ANDV N hinders autophosphorylation of TBK1, resulting in the inhibition of IRF3 phosphorylation and RIG-I/MDA5-directed type I IFN induction (36). Additionally, N can affect protein kinase R (PKR) dimerization (37), thereby preventing PKR phosphorylation, which is essential for its enzymatic activity. PKR inhibits virus replication (38).

Bunyaviral NSs are nonessential for virus replication, but they are pathogenesis determinants by acting as IFN antagonists (39). As a case in point, ANDV/CHI-9717869 NSs antagonize the type I IFN induction pathway (20). We therefore hypothesize that the two changes observed in NSs of ANDV strains associated with person-to-person transmission might enhance IFN antagonist potential. Moreover, the number of changes in ANDV/CHI-9717869 compared with ANDV/CHI-7913 and ANDV/ARG might explain the differences in lethality in the golden hamster animal model.

In the M segment, the amino-acid change T641I is also shared among ANDV strains associated with person-to-person-transmission but not among ANDV-like viruses. However, the change is also found in ANDV/NRC-6/18, which has not been associated with person-to-person transmission and it is absent in ANDV/NRC-3/18, which has been involved in an event of secondary transmission (**Table 3**). T641 is located in the signal peptide of Gc, in the region preceding the hyperconserved cleavage site WAASA. Because host protease landing sites are guided by the signal from this region (20), we hypothesize that this mutation might affect the dynamics and speed of ANDV glycoprotein retention and trafficking. Signal peptides share several characteristic features determined by their amino-acid composition (40), including a tripartite architecture with a positively charged N-terminus and a hydrophobic segment that determines the strength of the signal.

The phylogenetic analysis showed that ANDV/ARG is closely related to variants causing disease in humans and groups according to their geographic origin. ANDV/CHI-7913 is most closely related to ANDV/ARG, more than sequences obtained from patients reported in the endemic region. ANDV/CHI-9717869, on the other hand, is the most genetically divergent and remote geographically. Indeed, ANDV/ARG and ANDV/CHI-7913 share the most positions compared to ANDV/CHI-9717869. Thus, the decision to use ANDV/CHI-9717869 as the accepted challenge stock for medical countermeasure assessment needs to be revised, as this strain is a clear outlier that might not be representative of wild-type circulating strains.

Taken together, the results of our study indicate that determination and subsequent comparison of wild-type, cell-culture-passaged, and animal model-derived ANDV—and likely other orthohantavirus genome sequences—may allow predictions regarding their overall virulence and transmissibility, possibly informing countermeasure approaches. To strengthen such predictions, additional sequence information from yet-to-be-characterized ANDV strains and completion of genomic sequences of ANDV-like viruses is warranted.

## MATERIALS AND METHODS

### Viruses and cells

Andes virus strain ARG (ANDV/ARG) was isolated from a long-tailed colilargo (*Oligoryzomys longicaudatus* (Bennett, 1832)) in grivet (*Chlorocebus aethiops* (Linnaeus, 1758)) kidney epithelial Vero E6 cells (CRL-1586; ATCC, Manassas, VA, USA) (41). Continuous ANDV infection of cells was monitored by immunofluorescence performed with a rabbit polyclonal serum generated against ANDV nucleocapsid protein (N) open reading frame (ORF) and real-time reverse transcription PCR (RT-qPCR), and cultures were passaged blindly. Serial passaging (p9–p19) was performed at a multiplicity of infection of 0.1.

### Pathogenicity assessment

An established lethal animal model of ANDV infection, using golden hamsters (*Mesocricetus auratus* (Waterhouse, 1839)) (12), was leveraged to compare the previously established pathogenicity of the ANDV strain ARG (ANDV/ARG p9) (17) and to assess the pathogenicity of ANDV/ARG p19. Eight 12-week-old golden hamsters (four males and four females, obtained from the Instituto Nacional de Producción de Biológicos in Buenos Aires) were exposed intramuscularly to 100 μL of mock inoculum (phosphate-buffered saline [PBS]). Nine 12-week-old golden hamsters (four males and five females) were exposed intramuscularly to 100 μL of PBS containing 10^5^ focus-forming units of ANDV/ARG p19. Exposed golden hamsters were placed individually in ventilated cages and monitored daily up to 33 d post-exposure. Food and water were available *ad libitum*. All animal experiments were performed in an accredited animal biological safety level 3 (ABSL-3) biocontainment laboratory in compliance with institutional guidelines and Argentinian national law no. 14,346, which regulates experiments involving animals and adheres to principles stated in the Guide for the Care and Use of Laboratory Animals, National Research Council.

### RT-qPCR

Lung specimens were obtained from all golden hamsters following standard necropsy protocols. Total RNA was extracted from lung specimens using Trizol, as described previously (42). Quantitative RT-qPCR using ANDV genomic small (S) segment primers was performed following published procedures (43). Two microliters of each RNA sample were amplified in duplicate assays with a CFX detection system (Bio-Rad, Hercules, CA, USA) using TaqMan RT-PCR master mix (Quanta Biosciences, Gaithersburg, MD, USA), according to the manufacturers’ instructions. A primer set designed to detect the human RNaseP gene was used to ensure that samples were free of PCR inhibitors and that RNA extractions were homogeneous.

### Genomic and phylogenetic analyses

Virus genome sequencing was performed using three ANDV cell-culture passages (early [p3], intermediate [p9], and late [p19]; cryopreserved lung tissue from a naturally ANDV-infected long-tailed colilargo (p0); and lung tissues obtained from a golden hamsters exposed to ANDV/ARG p9. Also included in the analysis was a blood clot sample from a hantavirus pulmonary syndrome (HPS) patient (case C1-s, survivor, 14 yr) associated with secondary transmission of Buenos Aires virus (BASV) in Central Argentina (24).

Total RNA was extracted from cell-culture supernatants, lung tissues, and clinical samples utilizing Trizol. Virus genome sequencing was performed as previously described (11, 44). Briefly, a targeted bait-enrichment approach was used to enrich RNA-seq libraries for sequencing on the MiSeq platform (Illumina, San Diego, CA, USA). Hantavirus sequences from each genomic segment (S, M, and L) were collected (Table S1) and aligned using MAFFT v.7.397, implemented in Clustal W version 2.0 (45). The initial dataset consisted of coding-complete sequences obtained in this work and listed in Table S1. Other American orthohantavirus sequences from GenBank were also included. The resulting alignments were visually inspected to identify synonymous and nonsynonymous changes. Phylogenetic trees were reconstructed using IQ-TREE v. 1.6.12 (46) with automatic model selection (47). Branch supports were assessed by 1,000 ultrafast bootstraps (46).

## Data availability

Sequencing data are publicly available through GenBank under accession numbers

OP555720 to OPG555735.

## ACKNOWLEDGMENTS

We gratefully acknowledge Silvia A. Girard for her excellent technical assistance, M. Amoroso for veterinary medical care, and Alexis Edelstein for access to the animal biological safety level 3 facilities. The authors also thank Anya Crane (Integrated Research Facility at Fort Detrick, National Institute of Allergy and Infectious Diseases, National Institutes of Health, Fort Detrick, Frederick, MD, USA) for critically editing the manuscript and Jiro Wada (Integrated Research Facility at Fort Detrick, National Institute of Allergy and Infectious Diseases, National Institutes of Health, Fort Detrick, Frederick, MD, USA) for preparing figures.

This work was supported in part through Laulima Government Solutions, LLC, prime contract with the National Institutes of Health (NIH) National Institute of Allergy and Infectious Diseases (NIAID) under Contract No. HHSN272201800013C. J.H.K. performed this work as an employee of Tunnell Government Services (TGS), a subcontractor of Laulima Government Solutions, LLC, under Contract No. HHSN272201800013C. The views and conclusions contained in this document are those of the authors and should not be interpreted as necessarily representing the official policies, either expressed or implied, of the U.S. Department of Health and Human Services, the U.S. Army, or of the institutions and companies affiliated with the authors, nor does mention of trade names, commercial products, or organizations imply endorsement by the U.S. Government.

We declare no conflict of interest.

---

**Supplementary Figure 1. Phylogenetic analysis of the M and L segments informs the evolutionary history of Buenos Aires virus (BASV) and Andes virus strain ARG (ANDV/ARG).**

All variants are listed with the strain name, region of origin, year of the isolation and accession number. Different colors are used for identification (brown for non-ANDV South American orthohantaviruses; green for ANDV-like viruses; light blue for ANDV strains; and dark blue for passaged strains in clade 3). Detailed information of epidemiological history of the strains is listed in **Table S1**.

**Table S1.** List of sequences and strains utilized in the comparative genomics study of American orthohantaviruses.

| Orthohantavirus | Source (human, rodent species) | Strain name | Clade (based on S segment) | Site/Source | Administrative region | Country | Year | Genbank accession number |  |  | Reference | PMID | Denomination in manuscript |
| --- | --- | --- | --- | --- | --- | --- | --- | --- | --- | --- | --- | --- | --- |
|  |  |  |  |  |  |  |  | S | M | L |  |  |  |
| Andes virus | Human | NRC-4/18 | 1 | Villa Meliquina | Neuquén | Argentina | 2018 | MN258226 | MN258192 | MN258159 | Martinez VP *et al.* | 33264545 | ANDV/NRC-4/18 |
| Andes virus | long-tailed colilargo (*Oligoryzomys longicaudatus*) | ARG | 3 | San Martín de los Andes | Neuquén | Argentina | 1999 | OP555728 | OP555724 | NA | **This work** |  | ANDV/ARG |
| Andes virus | Cell culture passage 3; Vero E6 | ARG/p3 | 3 | San Martín de los Andes | Neuquén | Argentina | 1999 | OP555720 | OP555725 | OP555732 | **This work** |  | ANDV/ARG p3 |
| Andes virus | Cell culture passage 9; Vero E6 | ARG/p9 | 3 | San Martín de los Andes | Neuquén | Argentina | 1999 | OP555721 | OP555726 | OP555733 | **This work** |  | ANDV/ARG p9 |
| Andes virus | Cell culture passage 19; Vero E6 | ARG/p19 | 3 | San Martín de los Andes | Neuquén | Argentina | 1999 | OP555722 | OP555727 | OP555734 | **This work** |  | ANDV/ARG p19 |
| Andes virus | ANDV/Arg p9-infected golden hamsters | Mau /ARG/p19 | 3 | San Martín de los Andes | Neuquén | Argentina | 1999 | OP555723 | OP555729 | OP555735 | **This work** |  | ANDV/Mau /ARG/p19 |
| Andes virus | Human | ARG-Epuyen | 2 | Epuyén | Chubut | Argentina | 2018 | MN258239 | MN258205 | MN258172 | Martinez VP *et al.* | 33264545 | ANDV/Epuyen/18-19 |
| Andes virus | Human | NRC-6/18 | 2 | El Hoyo | Chubut | Argentina | 2018 | MN258228 | MN258194 | MN258161 | Martinez VP *et al.* | 33264545 | ANDV/NRC-6/18 |
| Andes virus | Human | ARG-Epilink | 2 | El Bolsón | Rio Negro | Argentina | 1996 | MN258223 | MN258189 | MN258156 | Martinez VP *et al.* | 33264545 | ANDV/Epilink/96 |
| Andes virus | Human | NRC-2/97 | 2 | Bariloche | Rio Negro | Argentina | 1997 | MN258224 | MN258190 | MN258157 | Martinez VP *et al.* | 33264545 | ANDV/NRC-2/18 |
| Andes virus | Human | AREB14/P2 | 2 | El Bolsón | Rio Negro | Argentina | 2014 | MN850084 | MN850089 | MN850094 | Martinez VP *et al.* | 33264545 | ANDV/AREB14-P2/Hum/RioNegro-ARG/2014 |
| Andes virus | long-tailed colilargo (*Oligoryzomys longicaudatus*) | CHI-9717869 | 4 | Coyhaique | Aysén | Chile | 1997 | AF291702 | AF291703 | AF291704 | Meissner *et al.* | 12367756 | ANDV/CHI-9717869 |
| Andes virus | long-tailed colilargo (*Oligoryzomys longicaudatus*) | CHI-9717869 | 4 | Coyhaique | Aysén | Chile | 1997 | MT956622 | MT956623 | MT956621 | Warner *et al.* | 33627395 | ANDV/CHI-9717869 |
| Andes virus | Human | CHI-7913 | 1 | Mulchén | Biobio | Chile | 1999 | MT956618 | MT956619 | MT956620 | Warner *et al.* | 33627395 | ANDV/CHI-7913 |
| Andes virus | Human | CHI-7913 | 1 | Mulchén | Biobio | Chile | 1999 | AY228237 | AY228238 | AY228239 | Tischler *et al.* | 14513715 | ANDV/CHI-7913 |
| Buenos Aires virus | Human | Hu39694 | ANDV-like | Pergamino | Buenos Aires | Argentina | 2002 | AF482711 | NA | NA | Bohlman *et al.* | 11907216 | BASV/Hu39694 |
| Buenos Aires virus | Human | Hu39694 | ANDV-like | Pergamino | Buenos Aires | Argentina | 2002 | NA | AF028023 | NA | Levis *et al.* | 9498428 | BASV/Hu39694 |
| Buenos Aires virus | Human | BA02-C15 | ANDV-like | La Plata | Buenos Aires | Argentina | 2002 | OP555730 | OP555731 | NA | **This work** |  | BASV/BA02-C15 |
| Orán virus | long-tailed colilargo (*Oligoryzomys longicaudatus*) | OI22996 | ANDV-like | Orán | Salta | Argentina | 2002 | AF482715 | AF028024 | NA | Levis *et al.* | 9498428 | ORNV/OI22996 |
| Orán virus | Chacoan colilargo (*Oligoryzomys chacoensis*) | AND Nort | ANDV-like | Orán | Salta | Argentina | 2002 | AF325966 | NA | NA | Gonzalez Della Valle *et al.* | 12224579 | ORNV/AND Nort |
| Lechiguanas virus | Fluorescent colilargo (*Oligoryzomys flavescens*) | 22819 | ANDV-like | Lechiguanas islands | Entre Rios | Argentina | 2002 | AF482714 | AF028022 | NA | Bohlman *et al.* & Levis *et al.* | 11907216 / 9100632 | LECIV/22819 |
| Maporal virus | Fulvous colilargo (*Oligoryzomys fulvescens*) | HV-97021050 | AH | Western Venezuela | Western Venezuela | Venezuela | 2004 | AY267347 | AY363179 | EU788002 | Fulhorst *et al.* | 15246651 | MAPV/HV-97021050 |
| Rio Mamoré virus | Small-eared colilargo (*Oligoryzomys microtis*) | HTN-007 | AH | Iquitos | Maynas | Peru | 2010 | FJ532244 | FJ608550 | FJ809772 | Richter *et al.* | 20687859 | RIOMV/HTN-007 |
| Laguna Negra virus | Little laucha (*Calomys laucha*) | 510B | AH | Chaco | Chaco | Paraguay | 1997 | AF005727 | AF005728 | NA | Johnson *et al.* | 9375015 | LANV/510B |
| Laguna Negra virus | Little laucha (*Calomys laucha*) | H731172/BRA259 | AH | Nova Olimpia | Paraná | Brazil | 2007 | NA | NA | JX443696 | Firth *et al.* | 23055565 | LANV/H731172/BRA259 |
| Sin Nombre virus | Human | NM H10 | AH | Four Corners Area | New Mexico | USA | 1994 | L25784 | L25783 | L37901 | Spiropoulou *et al.* | 8178455/7494336 | SNV/NM H10 |
| Choclo virus | Fulvous colilargo (*Oligoryzomys fulvescens*) | 588 | AH | Panamá | Panamá | Panamá | 2015 | KT983771 | KT983772 | EF397003 | Cajimat/Kho *et al.* | Unpublished | CHOV/588 |

non-ANDV American orthohantavirus
\* as reported, but there are no reports of long-tailed colilargos in northern Argentina

## REFERENCES

1. Kuhn JH, Charrel RN. 2018. Arthropod-borne and rodent-borne virus infections, p 1489–1509. *In* Jameson JL, Fauci AS, Kasper DL, Hauser SL, Longo DL, Loscalzo J (ed), Harrison’s Principles of Internal Medicine, 20th ed, vol 2. McGraw-Hill Education, Columbus, USA.

2. Alonso DO, Iglesias A, Coelho R, Periolo N, Bruno A, Córdoba MT, Filomarino N, Quipildor M, Biondo E, Fortunato E, Bellomo C, Martínez VP. 2019. Epidemiological description, case-fatality rate, and trends of hantavirus pulmonary syndrome: 9 years of surveillance in Argentina. J Med Virol 91: 1173–1181.

3. Martinez VP, Bellomo CM, Cacace ML, Suárez P, Bogni L, Padula PJ. 2010. Hantavirus pulmonary syndrome in Argentina, 1995-2008. Emerg Infect Dis 16:1853–60.

4. Wells RM, Sosa Estani S, Yadon ZE, Enria D, Padula P, Pini N, Mills JN, Peters CJ, Segura EL, Hantavirus Pulmonary Syndrome Study Group for Patagaonia. 1997. An unusual hantavirus outbreak in southern Argentina: person-to-person transmission? Emerg Infect Dis 3:171–4.

5. Tortosa F, Carrasco G, Gallardo D, Prandi D, Parodi V, Santamaría G, Ragusa M, Volij C, Izcovich A. 2022. Factores pronósticos de síndrome cardiopulmonar y muerte por hantavirus Andes Sur: estudio de cohorte en San Carlos de Bariloche y su zona de influencia sanitaria. Medicina (B Aires) 82:351–360.

6. Levis S, Morzunov SP, Rowe JE, Enria D, Pini N, Calderon G, Sabattini M, St Jeor SC. 1998. Genetic diversity and epidemiology of hantaviruses in Argentina. J Infect Dis 177:529–38.

7. Jonsson CB, Schmaljohn CS. 2001. Replication of hantaviruses. Curr Top Microbiol Immunol 256:15–32.

8. Padula PJ, Edelstein A, Miguel SDL, López NM, Rossi CM, Rabinovich RD. 1998. Brote epidémico del síndrome pulmonar por hantavirus en la Argentina. Evidencia molecular de la transmisión persona a persona del virus Andes. Medicina (B Aires) 58 Suppl 1:27–36.

9. Enria D, Padula P, Segura EL, Pini N, Edelstein A, Posse CR, Weissenbacher MC. 1996. Hantavirus pulmonary syndrome in Argentina. Possibility of person to person transmission. Medicina (B Aires) 56:709–11.

10. Riquelme R, Rioseco ML, Bastidas L, Trincado D, Riquelme M, Loyola H, Valdivieso F. 2015. Hantavirus pulmonary syndrome, Southern Chile, 1995-2012. Emerg Infect Dis 21:562–8.

11. Martínez VP, Di Paola N, Alonso DO, Pérez-Sautu U, Bellomo CM, Iglesias AA, Coelho RM, López B, Periolo N, Larson PA, Nagle ER, Chitty JA, Pratt CB, Díaz J, Cisterna D, Campos J, Sharma H, Dighero-Kemp B, Biondo E, Lewis L, Anselmo C, Olivera CP, Pontoriero F, Lavarra E, Kuhn JH, Strella T, Edelstein A, Burgos MI, Kaler M, Rubinstein A, Kugelman JR, Sanchez-Lockhart M, Perandones C, Palacios G. 2020. “Super-spreaders” and person-to-person transmission of Andes virus in Argentina. N Engl J Med 383:2230–2241.

12. Hooper JW, Larsen T, Custer DM, Schmaljohn CS. 2001. A lethal disease model for hantavirus pulmonary syndrome. Virology 289:6–14.

13. Milazzo ML, Eyzaguirre EJ, Molina CP, Fulhorst CF. 2002. Maporal viral infection in the Syrian golden hamster: a model of hantavirus pulmonary syndrome. J Infect Dis 186:1390–5.

14. Martinez VP, Padula PJ. 2012. Induction of protective immunity in a Syrian hamster model against a cytopathogenic strain of Andes virus. J Med Virol 84:87–95.

15. Toro J, Vega JD, Khan AS, Mills JN, Padula P, Terry W, Yadón Z, Valderrama R, Ellis BA, Pavletic C, Cerda R, Zaki S, Shieh W-J, Meyer R, Tapia M, Mansilla C, Baro M, Vergara JA, Concha M, Calderon G, Enria D, Peters CJ, Ksiazek TG. 1998. An outbreak of hantavirus pulmonary syndrome, Chile, 1997. Emerg Infect Dis 4:687–94.

16. Galeno H, Mora J, Villagra E, Fernandez J, Hernandez J, Mertz GJ, Ramirez E. 2002. First human isolate of hantavirus *(Andes virus)* in the Americas. Emerg Infect Dis 8:657–61.

17. Warner BM, Sloan A, Deschambault Y, Dowhanik S, Tierney K, Audet J, Liu G, Stein DR, Lung O, Buchanan C, Sroga P, Griffin BD, Siragam V, Frost KL, Booth S, Banadyga L, Saturday G, Scott D, Kobasa D, Safronetz D. 2021. Differential pathogenesis between Andes virus strains CHI-7913 and Chile-9717869 in Syrian Hamsters. J Virol 95:e00108–21.

18. Safronetz D, Prescott J, Feldmann F, Haddock E, Rosenke R, Okumura A, Brining D, Dahlstrom E, Porcella SF, Ebihara H, Scott DP, Hjelle B, Feldmann H. 2014. Pathophysiology of hantavirus pulmonary syndrome in rhesus macaques. Proc Natl Acad Sci U S A 111:7114–9.

19. Nemirov K, Lundkvist Å, Vaheri A, Plyusnin A. 2003. Adaptation of Puumala hantavirus to cell culture is associated with point mutations in the coding region of the L segment and in the noncoding regions of the S segment. J Virol 77:8793–800.

20. Vera-Otarola J, Solis L, Lowy F, Olguín V, Angulo J, Pino K, Tischler ND, Otth C, Padula P, López-Lastra M. 2020. The Andes orthohantavirus NSs protein antagonizes the type I interferon response by inhibiting MAVS signaling. J Virol 94:e00454–20.

21. Simons MJ, Gorbunova EE, Mackow ER. 2019. Unique interferon pathway regulation by the Andes virus nucleocapsid protein Is conferred by phosphorylation of serine 386. J Virol 93:e00338–19.

22. Duehr J, McMahon M, Williamson B, Amanat F, Durbin A, Hawman DW, Noack D, Uhl S, Tan GS, Feldmann H, Krammer F. 2020. Neutralizing monoclonal antibodies against the Gn and the Gc of the Andes virus glycoprotein spike complex protect from virus challenge in a preclinical hamster model. mBio 11:e00028–20.

23. Iglesias AA, Bellomo CM, Martínez VP. 2016. Síndrome pulmonar por hantavirus en Buenos Aires, 2009-2014. Medicina (B Aires) 76:1–9.

24. Martinez VP, Bellomo C, San Juan J, Pinna D, Forlenza R, Elder M, Padula PJ. 2005. Person-to-person transmission of Andes virus. Emerg Infect Dis 11:1848–53.

25. Löber C, Anheier B, Lindow S, Klenk H-D, Feldmann H. 2001. The Hantaan virus glycoprotein precursor is cleaved at the conserved pentapeptide WAASA. Virology 289:224–9.

26. von Heijne G. 1990. The signal peptide. J Membr Biol 115:195–201.

27. Koehler A, Kolesnikova L, Becker S. 2016. An active site mutation increases the polymerase activity of the guinea pig-lethal Marburg virus. J Gen Virol 97:2494–2500.

28. Trefry JC, Wollen SE, Nasar F, Shamblin JD, Kern SJ, Bearss JJ, Jefferson MA, Chance TB, Kugelman JR, Ladner JT, Honko AN, Kobs DJ, Wending MQS, Sabourin CL, Pratt WD, Palacios GF, Pitt MLM. 2015. Ebola virus infections in nonhuman primates are temporally influenced by glycoprotein poly-U editing site populations in the exposure material. Viruses 7:6739–54.

29. Boudko SP, Kuhn RJ, Rossmann MG. 2007. The coiled-coil domain structure of the Sin Nombre virus nucleocapsid protein. J Mol Biol 366:1538–44.

30. Wang Y, Boudreaux DM, Estrada DF, Egan CW, St Jeor SC, De Guzman RN. 2008. NMR structure of the N-terminal coiled coil domain of the Andes hantavirus nucleocapsid protein. J Biol Chem 283:28297–304.

31. Guo Y, Wang W, Sun Y, Ma C, Wang X, Wang X, Liu P, Shen S, Li B, Lin J, Deng F, Wang H, Lou Z. 2016. Crystal structure of the core region of hantavirus nucleocapsid protein reveals the mechanism for ribonucleoprotein complex formation. J Virol 90:1048–61.

32. Yoshimatsu K, Arikawa J. 2014. Antigenic properties of N protein of hantavirus. Viruses 6:3097–109.

33. Kanerva M, Melén K, Vaheri A, Julkunen I. 1996. Inhibition of Puumala and Tula hantaviruses in Vero cells by MxA protein. Virology 224:55–62.

34. Pavlovic J, Schröder A, Blank A, Pitossi F, Staeheli P. 1993. Mx proteins: GTPases involved in the interferon-induced antiviral state. Ciba Found Symp 176:233–43; discussion 243-7.

35. Khaiboullina SF, Rizvanov AA, Lombardi VC, Morzunov SP, Reis HJ, Palotás A, St Jeor S. 2013. Andes-virus-induced cytokine storm is partially suppressed by ribavirin. Antivir Ther 18:575–84.

36. Cimica V, Dalrymple NA, Roth E, Nasonov A, Mackow ER. 2014. An innate immunity-regulating virulence determinant is uniquely encoded by the Andes virus nucleocapsid protein. mBio 5:e01088–13.

37. Wang Z, Mir MA. 2015. Andes virus nucleocapsid protein interrupts protein kinase R dimerization to counteract host interference in viral protein synthesis. J Virol 89:1628–39.

38. Goodbourn S, Didcock L, Randall RE. 2000. Interferons: cell signalling, immune modulation, antiviral response and virus countermeasures. J Gen Virol 81:2341–2364.

39. Ly HJ, Ikegami T. 2016. Rift Valley fever virus NSs protein functions and the similarity to other bunyavirus NSs proteins. Virol J 13:118.

40. von Heijne G. 1999. Recent advances in the understanding of membrane protein assembly and structure. Q Rev Biophys 32:285–307.

41. Padula PJ, Sanchez AJ, Edelstein A, Nichol ST. 2002. Complete nucleotide sequence of the M RNA segment of Andes virus and analysis of the variability of the termini of the virus S, M and L RNA segments. J Gen Virol 83:2117–2122.

42. Padula P, Figueroa R, Navarrete M, Pizarro E, Cadiz R, Bellomo C, Jofre C, Zaror L, Rodriguez E, Murúa R. 2004. Transmission study of Andes hantavirus infection in wild sigmodontine rodents. J Virol 78:11972–9.

43. Bellomo CM, Pires-Marczeski FC, Padula PJ. 2015. Viral load of patients with hantavirus pulmonary syndrome in Argentina. J Med Virol 87:1823–30.

44. Alonso DO, Pérez-Sautu U, Bellomo CM, Prieto K, Iglesias A, Coelho R, Periolo N, Domenech I, Talmon G, Hansen R, Palacios G, Martinez VP. 2020. Person-to-person transmission of Andes virus in hantavirus pulmonary syndrome, Argentina, 2014. Emerg Infect Dis 26:756–759.

45. Katoh K, Standley DM. 2013. MAFFT multiple sequence alignment software version 7: improvements in performance and usability. Mol Biol Evol 30:772–80.

46. Minh BQ, Schmidt HA, Chernomor O, Schrempf D, Woodhams MD, von Haeseler A, Lanfear R. 2020. IQ-TREE 2: new models and efficient methods for phylogenetic inference in the genomic era. Mol Biol Evol 37:1530–1534.

47. Kalyaanamoorthy S, Minh BQ, Wong TKF, von Haeseler A, Jermiin LS. 2017. ModelFinder: fast model selection for accurate phylogenetic estimates. Nat Methods 14:587–589.

